# Predictive Modeling of Evoked Intracranial EEG Response to Medial Temporal Lobe Stimulation in Patients with Epilepsy

**DOI:** 10.1101/2023.08.07.552297

**Authors:** Gagan Acharya, Kathryn A. Davis, Erfan Nozari

## Abstract

Despite promising advancements, closed-loop neurostimulation for drug-resistant epilepsy (DRE) still relies on manual tuning and produces variable outcomes, while automated predictable algorithms remain an aspiration. As a fundamental step towards addressing this gap, here we study predictive dynamical models of human intracranial EEG (iEEG) response under parametrically rich neurostimulation. Using data from n = 13 DRE patients, we find that stimulation-triggered switched-linear models with ∼300ms of causal historical dependence best explain evoked iEEG dynamics. These models are highly consistent across different stimulation amplitudes and frequencies, allowing for learning a generalizable model from abundant STIM OFF and limited STIM ON data. Further, evoked iEEG in nearly all subjects exhibited a distance-dependent pattern, whereby stimulation *directly* impacts the actuation site and nearby regions (≲ 20mm), affects medium-distance regions (20 ∼ 100mm) through network interactions, and hardly reaches more distal areas (≳ 100mm). Peak network interaction occurs at 60∼80mm from the stimulation site. Due to their predictive accuracy and mechanistic interpretability, these models hold significant potential for model-based seizure forecasting and closed-loop neurostimulation design.

## 1 Introduction

Closed-loop neurostimulation has long been considered a promising alternative to highly invasive resection surgery for individuals with drug-resistant epilepsy. Extensive research over the past decades has highlighted its numerous advantages, including short-term benefits like electric stimulation-induced desynchronization of neuron firing to counteract seizures, as well as long-term benefits such as the modification of cortical excitability via neuroplasticity^1, 2^. Nevertheless, its state-of-the-art clinical implementation, the RNS^®^system from NeuroPace, Inc.^1^ has known limitations, including a lack of mechanistic understanding^3^, simplistic detection algorithms, and non-adaptive pre-selected stimulation parameters^1, 4, 5^. As a result, though highly effective in some individuals, RNS outcomes vary widely among patients, and only 18% of those receiving RNS implantation experience seizure freedom for a period of 1 year or longer^6^. To establish neurostimulation as a robust treatment with predictable outcomes, it is vital to have *the ability to predict* the future trajectory of neural activity, both in the absence and presence of neurostimulation. Such predictive modeling, in particular, opens the door to a host of model-based detection and control algorithms that have been developed and studied for decades in engineering^7–9^ but often need accurate predictive models to succeed.

Despite decades of research, accurate predictive models of an epileptic brain’s response to neurostimulation are largely lacking. A large group of studies have developed models that primarily provide explanations for the disorder on a phenomenological level^10–13^. These models have been used for extracting features in seizure detection algorithms^14–21^, conducting stability analysis^8, 22^, and localizing seizures^23^. However, these models are not tuned using subject-specific stimulation data and are thus inadequate for robust and predictable closed-loop control design over the diverse set of parameters for neurostimulation^8, 9^. A second body of work has pursued more accurate and/or subject-specific models using detailed finite-element modeling of the brain’s electromagnetic response to neurostimulation^24–27^. This is often done in conjunction with biophysical modeling and used for the analysis and optimization of the impact of electrode placement and (de)polarization activity of neuronal tissue. Nevertheless, these models are often not built for and capable of accurate time-series prediction, particularly at a large-scale network level beyond the local stimulation site.

Finally, a third body of work has specifically pursued data-driven predictive modeling of the brain’s response to neurostimulation. Multiple studies have used linear autoregressive models to fit the neural activity triggered by neurostimulation^28–31^ while others have opted for nonlinear kernel-based autoregressive modeling^32, 33^. However, these works have mainly focused on univariate modeling (i.e., modeling the neurostimulation-evoked response at a single measurement source), while modeling the evoked *network response* has remained more challenging^34–36^. Furthermore, a significant portion of these studies relies on simulated data, data from non-human primates, or data in the absence of stimulation. As such, accurate subject-specific predictive models of brain network dynamics under neurostimulation have remained largely lacking. Finally, several works have used cortico-cortical evoked potentials (CCEPs, a particular form of evoked iEEG response to intracranial stimulation) to study effective brain connectivity^37–39^. These works, however, differ from the present study in that they do not pursue dynamical system modeling of the evoked response, as pursued herein.

In this study, we present a rigorous data-driven framework for developing subject-specific network dynamical models of intracranial electroencephalography (iEEG) dynamics in the absence and presence of medial temporal lobe electrical stimulation in patients with epilepsy. We use recordings from the Restoring Active Memory (RAM) Public dataset, which offers a particularly rich resource for datadriven modeling due to the presence of factorial parameter search experiments where multiple stimulation parameters are varied in extensive combinations. We follow a standard system identification approach^40^ where models are compared based on the accuracy of their one-step-ahead predictions (namely, predictions of iEEG measurements at any sample *t* based on all the available histories of past iEEG and stimulation data up to sample *t* − 1). This emphasis on predictive modeling is not only essential for closed-loop seizure prediction and suppression, as noted earlier, but also provides the means for *studying Granger causality* among the input (neurostimulation) and output (iEEG) variables, as described next.

## 2 Results

### Model

The ability of any computational model to explain data depends heavily on its chosen structure. In this work, unless when otherwise stated, we use a bilinear model of the form

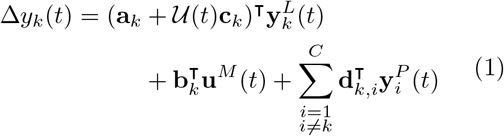

to model individualized (subject and channel-specific) iEEG response to stimulation. In this model, the one-step change in the iEEG activity of the *k*^*th*^ channel, i.e., Δ*y*_*k*_(*t*) = *y*_*k*_(*t* + 1) − *y*_*k*_(*t*), is predicted linearly based on the past *L* samples 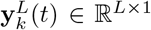 of iEEG in that same channel, the past *M* samples **u**^*M*^ (*t*) ∈ ℝ^*M ×*1^ of stimulation input, the past *P* samples 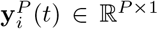 of iEEG in other channels, and the bilinear interactions 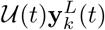 between the stimulation input and iEEG output (cf. Methods for details). While we focus on one-step ahead prediction, our results and methods can be extended to modeling the *k* −step ahead response as shown in Supplementary Figure 6. This bilinear choice of the model structure is motivated by the long history of bilinear modeling in neuroscience^41^ as well as our prior work showing the linearity of resting state (i.e., STIM OFF) iEEG^42^. We thus depart from linearity towards highly complex models (e.g., deep neural networks) gradually and only to the extent supported by data. Bilinear models provide a natural first step in moving beyond linearity^43–45^ and are thus preferred unless more complex models provide substantial improvement in accuracy, a topic to which we will return later (cf. Figure 6b).

### iEEG data contains about 300ms of causal historical dependence, despite significant heterogeneity across subjects and channels

Unlike models with latent states such as linear state-space models^40, 46^ or hidden Markov models^47^, autoregressive models predict future outputs based on the history (lags) of *observed* data, i.e., inputs and outputs. This creates a trade-off, where AR models have significantly larger state dimensions in return for having the state fully observed (and therefore known more accurately) and interpretable. A critical structural parameter for AR models is thus the amount of input and output history used for prediction. Notably, this is different from and can be significantly smaller than, the decay of iEEG autocorrelation. While the latter answers the question of whether the output at two separate time points is merely *correlated*, the former answers a *causal* question, in the Granger sense of causality^48^, of whether any piece of data in the past provides *additional information* beyond what is available in the less-distant past.

Figure 2a shows how autoregressive models with different amounts of autoregressive (*L*) and input (*M*) history compare across all subjects and channels. In general, the amount of historical dependence that gives the highest predictive accuracy greatly varies among subjects and even different channels within each subject. In fact, for almost all combinations of *L* and *M*, we can find at least one subject-channel where that combination is optimal. This vast heterogeneity exists regardless of the statistical measures for comparison (mean- or median-based), and reinforces the need for *sub-individualized* data-driven modeling, as opposed to normative models which explain seizure dynamics generically. In what follows, we evaluate each model on the basis of its ‘win rate’, i.e., the percentage of subject-channels for which that model had the smallest cross-validated predictive accuracy.

**Figure 1:**
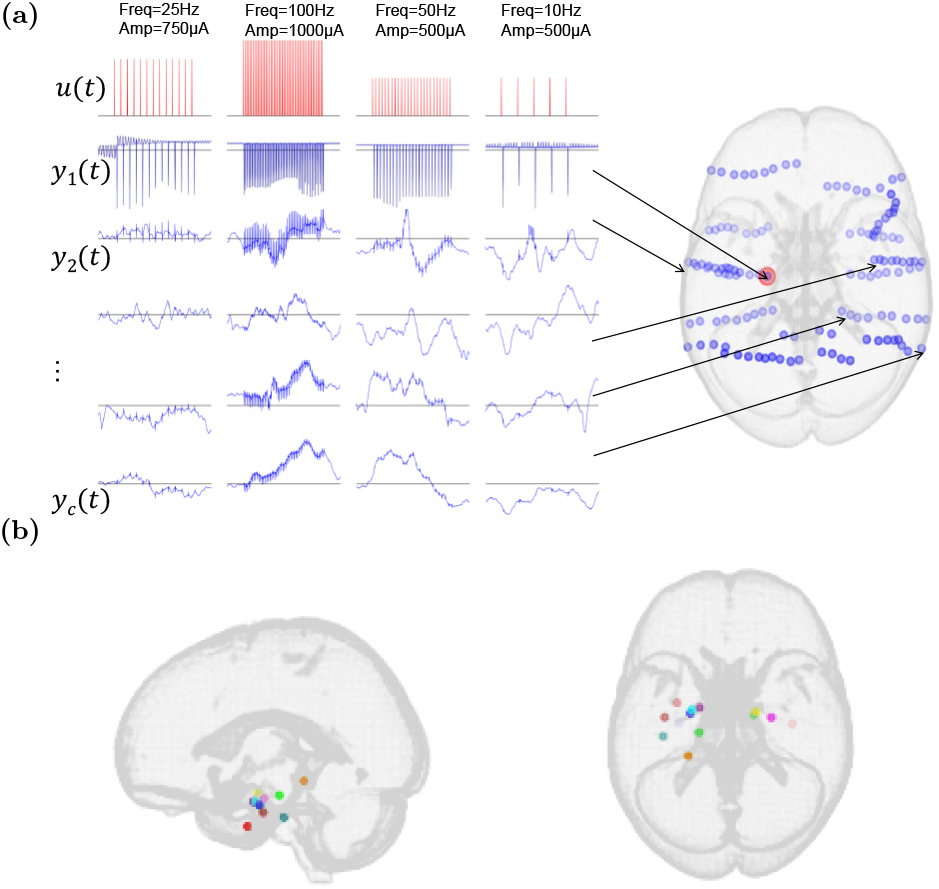
Illustration of the iEEG data used in this work. (a) Each cluster of adjacent blue circles corresponds to contacts on a single electrode (warped due to registration to standard space). *u*(*t*) encodes the applied stimulation pulse train, including both its frequency and amplitude. *y*_*k*_(*t*) refers to the recorded iEEG signal of the *k*^*th*^ electrode ordered based on its closeness to the stimulation site. (b) Stimulation locations across the 13 subjects in this study, shown in standard MNI space. Only subjects with Medial Temporal Lobe (MTL) stimulation were selected (see Methods for details).

**Figure 2:**
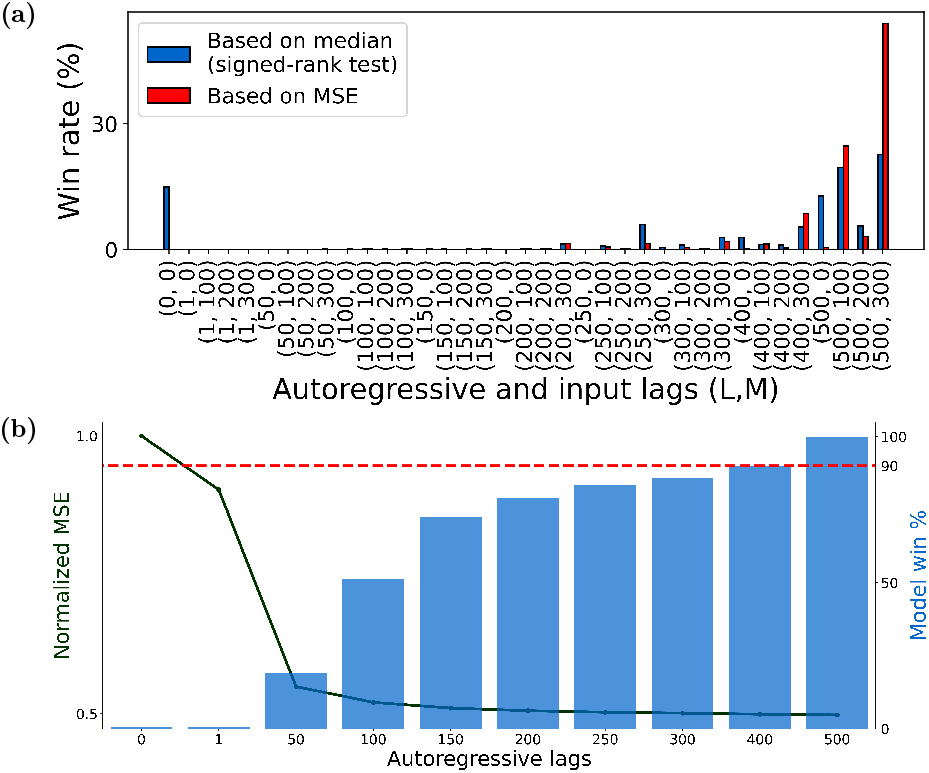
Model order distribution and performance across subjects and channels. For each subject-channel we trained ARX models of various orders (*L, M*) (*L* ∈ {0, 1, 50, 100, …, 500} and *M* ∈ {0, 100, 200, 300}) and determined the pair (*L, M*) that resulted in the model with the *best* fit over unseen data. (a) We plotted the *Win rate* i.e., the percentage of times each (*L, M*) pair is the *best* across all considered channels (*n* = 1547). We used 2 definitions of the *best* model: based on the mean ((*L, M*) pair that resulted in the lowest MSE) and based on the median ((*L, M*) pair that is superior to other model orders as per Wilcoxon signed-rank test (see Methods)). Irrespective of the definition used, we saw that larger autoregressive lags (i.e., *L* ≥ 400) are favored across more than 90% channels. About 66% of the modeled channels favored a non-zero input lag (i.e., *M* > 0) and hence are causally affected by direct stimulation input. (b) We also plotted the validation MSE (averaged across all *M*) as a function of the autoregressive lags (*L*) (left y-axis) and found that although a larger *L* is more favored, doing so only resulted in a marginally better fit in comparison to a model with 200 to 300 autoregressive lags. In the same figure, we also plot (right y-axis) the ‘statistical win rate’, defined as the percentage of channels where a model of order *L* has an MSE that is statistically similar (i.e., within ±2*S*.*E*.*M*) to the MSE corresponding to the best model. From this plot, we observed that a model with *L* = 300 is sufficient for nearly 90% of all the channels.

Increasing the amount of autoregressive history (*L*) almost monotonically improves predictive accurac but only marginally so beyond 200-300ms. Speci ically, models with larger *L* have larger win rate across subjects and channels (Figure 2b, bar plot but increasing autoregressive history from 300 t 500ms only improves win rate by 10% (Figure 2 dashed horizontal line). This trend can also be see from average prediction error (average normalize MSE across all subjects and channels), which drop significantly when increasing *L* from 1ms to 50ms an decays fairly notably until about 200-300ms but saturates thereafter (Figure 2b, solid line).

Unlike autoregressive history, the amount of past input history (*M*) that has a causal effect on evoked iEEG response varies greatly among subjects and channels (Figure 2a). For about two-thirds of the subject-channels, some (though still heterogeneous) amount of input history improves predictive accuracy, while for other subject-channels inclusion of any amount of input history, even from the immediate past, lowers predictive accuracy. This lack of direct causal effect in the latter group is consistent with the hypothesis that not all channels are directly excited by stimulation, but rather receive the impact of stimulation through complex spatiotemporal interactions with other channels. Such network-mediated stimulation effects may also be small enough to be statistically undetectable compared to baseline fluctuations^42^, characterizing a group of subject-channels with no evoked response, whether direct or indirect (see also Figure 5).

Given the relatively fast sampling rate of iEEG (1000Hz), it is also possible that successive input/output samples contain redundant, correlated information and need not all be included in the model. To test this hypothesis, we compared the above-described models with densely-sampled input-output history against sparsely-sampled alternatives where both the input and output are sampled every *τ* milliseconds (*τ* > 1). The same number of lags was used across the dense and sparsely sampled models to ensure fair comparisons. We find that at low model lags, sparse sampling is more beneficial indicating that the inherent process is more dependent on dynamics over longer time horizons as opposed to highly correlated dense samples. With increasing lags however (Figure 3), dense models achieve higher predictive accuracy, even though sparse models with the same number of lags have access to a wider window for prediction (*τ L*ms for a sparse model compared to *L*ms in a dense model). In particular, this result *discourages the use of delay embedding* ^49^ despite its theoretical appeal^50^, and instead encourages investing the available computational power (which is often extremely limited in chronic implants) into learning temporally short-range predictive relationships.

**Figure 3:**
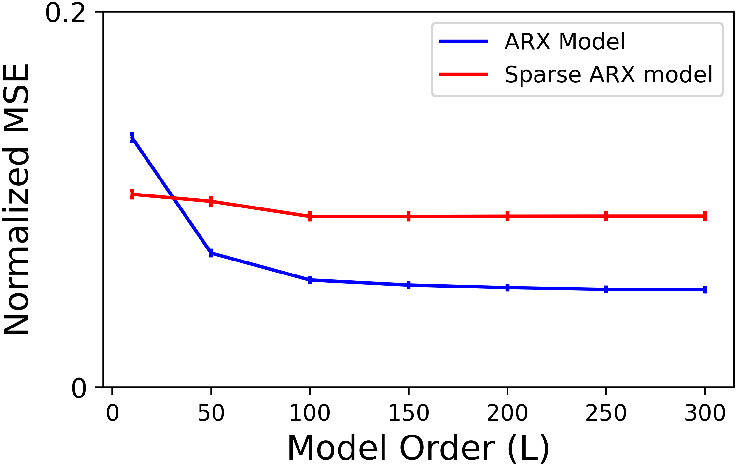
Effect of sparser temporal sampling of autoregressive lags on model accuracy. For each channel in the data, we considered 2 categories of models: ARX models with *dense temporal sampling* (i.e., iEEG lags are spaced 1ms apart) and ARX models with *sparse temporal sampling* (i.e., iEEG lags are spaced *τ* > 1ms apart). For the latter, the exact sampling time *τ* was chosen separately for each channel based on delay embedding theory using the method described in Otani et al.^51^. We trained sparse and dense models of various lags (*L* ∈ 1, 50, 100, 150, 200, 250, 300) and plotted the normalized MSE across subject-channels as a function of *L*. Although the sparse ARX model with *L* lags has access to a much larger duration of iEEG history (i.e., *τL*ms) in comparison to a dense model (i.e., *L*ms), the former has lower accuracy. Thus, temporally short-range models with higher sampling rates are more predictive.

### iEEG evoked response is best described by switched-linear dynamics

The functional form, extent, and even the very presence of nonlinearity in the neural dynamics of patients with epilepsy has been the topic of much debate^52, 53^. In our recent work^42^, we showed that resting state (interictal, STIM OFF) iEEG dynamics are best described by linear autoregressive models. However, our examination of STIM ON and STIM OFF periods reveals notable distinctions in the optimal autoregressive parameters (i.e., **a**_*k*_ in Eq. (1)) when fit separately for each duration. Therefore, we first sought direct evidence for whether the presence of stimulation makes iEEG dynamics nonlinear. We tested if parameter **a**_*k*_ trained using resting state data (i.e., STIM OFF) can generalize over STIM ON durations. We did this by comparing 2 linear models across each subject: (1) an ARX model trained over STIM ON duration (Stim Model in Figure 4a) and (2) the Stim Model with the **a**_*k*_ vector replaced with that trained using STIM OFF data (RS Model in Figure 4a). If the dynamics in the STIM ON and STIM OFF durations were indeed similar, we could expect the above 2 models to perform similarly when we test them on unseen data. Moreover, an agreement between these models would indicate that the iEEG dynamics can be captured largely by a single linear model. However, our analysis in Figure 4a and 4b showed that the Stim Model performs significantly better on unseen data than the RS Model and thus indicated the presence of stimulation-induced nonlinearity. Note that the lag parameters *L* and *M* for the above experiment, as well as for the remainder of the analysis in this work, were chosen based on the results shown in Figure 2. For each subject-channel, we used the optimal (*L, M*) pair, as indicated by the median-based performance. Arguably, the simplest nonlinear choice to account for the observed differences between STIM ON and STIM OFF dynamics is a switched-linear model that simply switches between one linear model during STIM ON and another linear model during STIM OFF. This is also a special case of the bilinear model in Eq. (1), wherein the gating term 𝒰(*t*) = 1 if *t* is within a stimulation duration and 0 otherwise. A direct comparison between ARX and switched ARX models showed that the switched ARX model was favored over its linear counterpart for the majority of subject-channels, as illustrated in Figure 4c.

**Figure 4:**
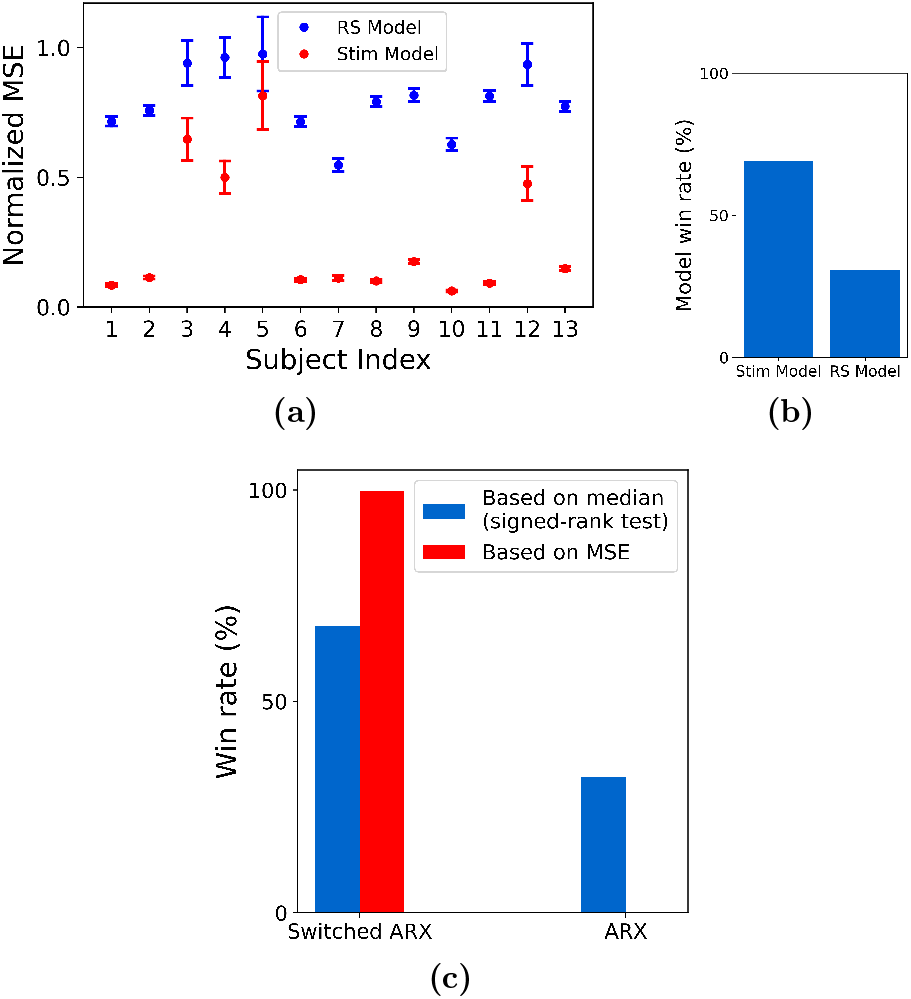
Evidence of nonlinear dynamics and switching behavior. We tested an ARX model trained using STIM ON (Stim model) data against a similar model with the **a**_*k*_ matrix replaced by that trained using STIM OFF data (Resting State (RS) model). (a) The performance (MSE) difference between these models and (b) their respective win rates, indicating the presence of stimulation-dependent nonlinearity. (c) When we account for the nonlinearity using a switched ARX model, we found it to be more descriptive of the iEEG response in comparison to a single ARX model

**Figure 5:**
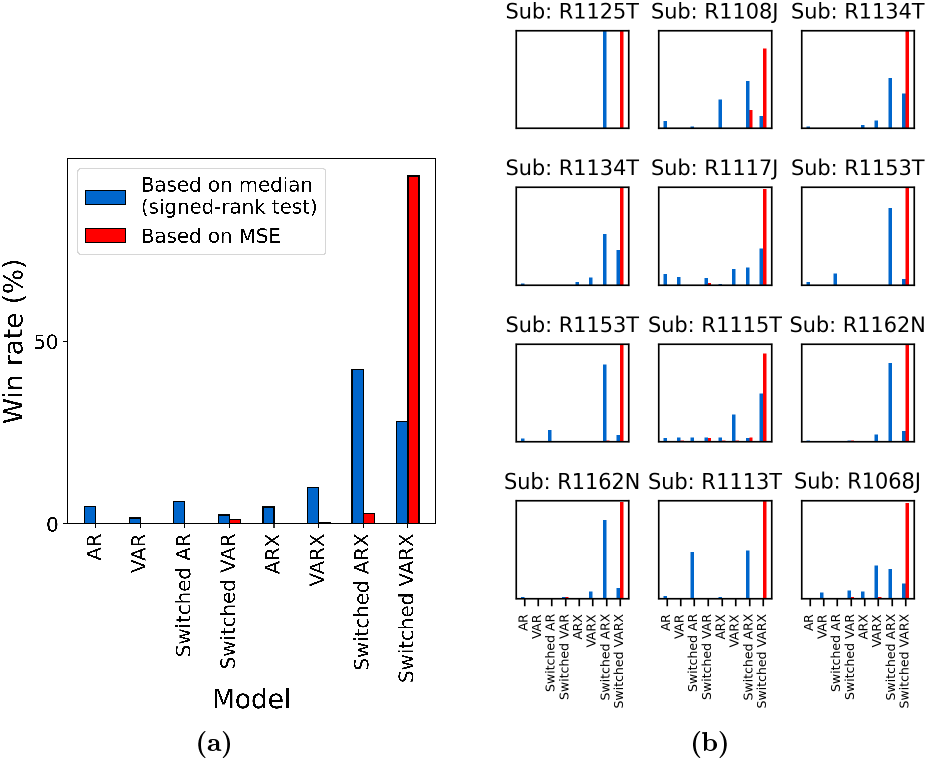
Comparison of linear and switched-linear models. (a) Win-rate distribution across all subjects and channels. Irrespective of the criterion used, switched-linear models were observed to outperform linear models for more than 90% of the channels. The preference for modeling the causal effects of stimulation and network interactions seemed to vary greatly among channels. (b) A zoomed-in version of the plot in (a) shows the subject-wise distributions of the model win rate.

**Figure 6:**
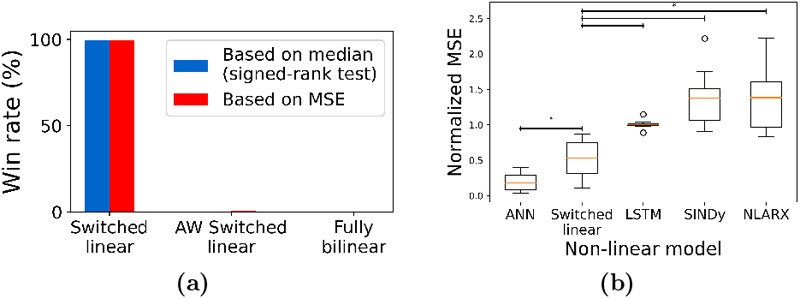
Comparison of switched-linear models with other forms of nonlinear models. (a) We compared the switched-linear model against an amplitude-weighted (AW) switched-linear model and a fully-bilinear model. As indicated by the win rate across, we found switched-linear models to be most predictive across all subject-channels (b) When we compared switched-linear to other forms of nonlinear models, we found that the artificial neural networks (ANNs) resulted in the lowest MSE and this result was followed by the switchedlinear model (\**p* < 0.05).

Nevertheless, the comparison of the descriptive ability of a switched-linear model to that of a simple linear model has to be made from a more nuanced level taking into account the potentially confounding effects of other model attributes, such as the presence of network interactions and direct STIM effect. Motivated by this, we compared a host of linear models, such as the VARX model (linear ARX model with network interactions) or AR model (linear AR model), with their switched-linear counterparts. As shown in Figure 5a, independent of the statistical measure for comparison (mean or median), the switched-linear class of models is more consistent with the data in the vast majority of cases. That being said, there is still vast heterogeneity among subjects and channels in what kind of switched-linear model achieves the highest predictive power, while even some (about 10%) of subject-channels are best described by a single linear model throughout all STIM ON and STIM OFF durations. The distributions of win rates for each subject also show a consistent preference for switched-linear models across all subjects, even though vast heterogeneity also persists in these distributions, both within and between subjects (Figure 5b).

A further aspect of the large heterogeneity in the channel models in Figure 5 is whether stimulation has a direct causal effect on iEEG evoked response. We observed that stimulation effects are predominant in more than 75% cases across subject-channels. This observation was further validated when we conducted a bootstrapping experiment (see Methods) over the input lags and found empirical evidence for direct stimulation effects in about 95% of the 75% subject-channels considered in the test.

Despite the higher predictive accuracy of switched-linear models compared to linear ones, their binary gating enforces a single linear model for all STIM ON durations, irrespective of the stimulation’s frequency and amplitude. A fully bilinear model with 𝒰(*t*) = *u*(*t*) in the form described by equation Eq. (1) can potentially alleviate this issue by allowing the stimulation waveform to directly modulate the linear dynamics using the interaction term 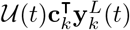. A third, middle-ground alternative is to employ an amplitude-weighted switched-linear (AWSL) model where the gating term 𝒰(*t*) is determined by the amplitude of the stimulation rather than a binary signal. However, we found the switched-linear model to be more consistent with the data compared to the more complex bilinear and AWSL alternatives (Figure 6). This is remarkable not only from a biological perspective but also from an engineering perspective given the large body of literature on control design for switched linear systems^54^.

While we notice the lack of benefit in bilinear over switched-linear models, this does not remove the possibility that significantly more complex models still explain stimulation-evoked iEEG response better. When compared against some of the most commonly known nonlinear models, switched-linear models were highly predictive across channels, second only to feedforward Artificial Neural Networks (ANNs) with *ReLU* activation. Note, also, that ANNs with *ReLU* activation are also switched-linear models. In fact, when compared against ANNs with *tanh* nonlinearity, those with *ReLU* activations explained the data significantly better. Nevertheless, compared to a simple switched-linear model in Eq. (1), the ANNs with *ReLU* activation have two orders of magnitude more parameters, combinatorially more switching regions, and almost no biological interpretability. This makes the former, simpler switched-linear model potentially preferred for mechanistic understanding and model-based closed-loop seizure stabilization, particularly in chronic implants with limited logic gates and battery resources.

### Network-mediated input spread is consistently distance-dependent

A critical feature in the dynamics of any iEEG channel is network interactions with other channels, both from a network control perspective as well as a mechanistic understanding of how stimulation effects spread throughout the brain. Network interactions are captured by the term 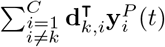 in Eq. (1), where we refer to models with at least one nonzero **d**_*k,i*_ (at least one nonzero effective connectivity between two electrodes) as *vector* autoregressive (VAR). In our recent work^42^, we found VAR models to be *less*, not more, predictive of resting state iEEG compared to models with no network interactions (all **d**_*k,i*_ = **0**), indicating a lack of causal effects among the channels at rest.

The presence of stimulation, on the other hand, has the potential to boost causal network interactions through which the stimulation effect can spread. As noted earlier (cf. Figure 2a), across all subjects, we found no *direct* causal effect from stimulation input to about 25% of the channels. Nevertheless, several of such channels still demonstrate clearly heightened activity during STIM ON periods, therefore hinting at an *indirect* causal effect of stimulation mediated by network interactions (Figure 7).

**Figure 7:**
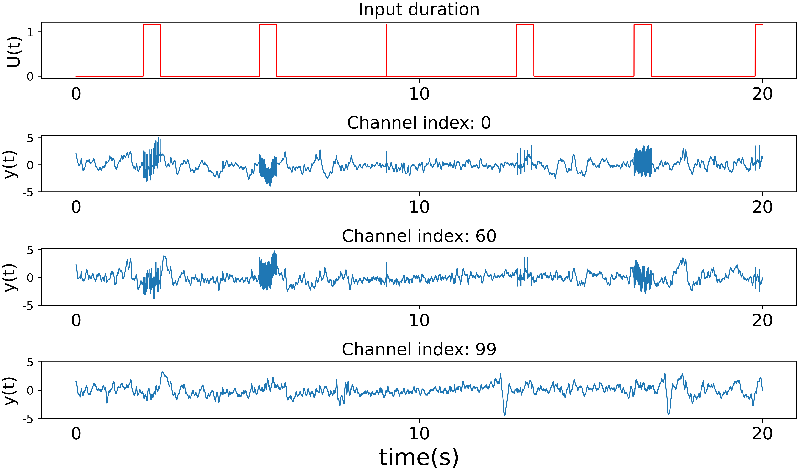
Heightened activity in channels without direct causal effect from the stimulation. (a) Plot showing the STIM ON durations within a specific stretch of the recordings. In (b)-(d), we plot the iEEG activity in response to the stimulation signal in (a) for 3 channels that did not show any evidence of being directly affected by the applied stimulation.

Assuming that the spread of the stimulation through the brain connectome is the main reason for the significance of network effects in STIM ON durations compared to resting state periods, then the stimulation channel itself should have no causal network effect *from* other channels. Therefore, we hypothesize that response at the site of stimulation can be best explained by a switched-linear model without network interactions (i.e., switched ARX). This was indeed the case when we compared switched VARX and switched ARX models trained for stimulation channels, and found that the latter model was favored in 10 out of the 13 subjects in our data. Even for the 3 subjects where switched VARX was favored, we found that the relative improvement in MSE from incorporating data from other channels was only modest (< 10%). Interestingly, this is consistent with the role played by the do-operator in causal discovery^55^, effectively isolating the anode from being causally influenced by other nodes within the network.

Analyzing the channels corresponding to electrodes that were directly stimulated, we consistently found that these non-stimulated channels most often receive causal effects from other nodes in the network. More interestingly, however, we found that the strength of this causal effect (i.e., the amount of added model predictive power when including the past history from other channels) varies in a consistently distant-dependent manner. At one extreme, we observed that the strength was least pronounced for electrodes located in close proximity to the anode. This indicates that the response in these channels, similar to the stimulated channel itself, is mostly overpowered by direct stimulation effects. On the other extreme are nodes that receive little to no effect from stimulation, whose network interactions are expectedly similar to weak/insignificant network interactions during resting state found earlier^42^. At an intermediary Euclidean distance from the anode (around 70±16 mm) were nodes that received the strongest causal effect from network interactions.

### Channel importance as a function of distance indicates an S-shaped relationship

An important inquiry in modeling dynamics across large networks pertains to whether precise predictions hinge on capturing network interactions across all channels or if comparable performance can be attained using only a limited subset of channels. In particular, does incorporating electrodes in close proximity to the channel being modeled suffice? If so, what is the radius of influence beyond which electrodes cease to significantly impact modeling (in the sense of Granger causality)?

We partially address this via backward elimination of regression features to compute the impact of removing network information pertaining to subsets of electrodes. Specifically for each subject-channel being modeled, we group electrodes based on their distance to the channel. Subsequently, for each group, we iteratively modeled the channel dynamics by eliminating the terms in Equation Eq. (1) associated with the electrodes in the group and compute the validation NMSE. We consider a channel to be successfully modeled if the obtained performance is *equivalent to* a model with complete network information (i.e., with all channels) as indicated by a < 2% relative increase in NMSE. As depicted in Figure 9, the graph of the percentage of successfully modeled channels against the distance of the eliminated electrode group demonstrates an S-shaped pattern (*p* < 0.001), suggesting that electrodes in close proximity exert a more pronounced influence on modeling outcomes.

The modeling error is the highest when removing local channels, as compared to those mid-distance from the target channel (Supplementary Figure 5). Specifically, we find that for approximately 50±12% of the channels, it is sufficient to include just the local electrodes within a 40mm radius of the modeled channel. This proportion rises to 80±9% of the channels when the radius is increased to 60mm. These findings hint at the potential for modeling iEEG response on a local scale and the prospect of constructing models with fewer regression features. Such models, as proposed, could be better suited for designing stimulation protocols in chronic implants.

### Evoked iEEG dynamics are highly consistent across stimulation frequencies but less consistent across sessions

A critical challenge for model-based control design for neurostimulation is learning models that generalize beyond the (past) data over which they are trained, particularly given the extreme inter and intra-subject variability in epilepsy on the one hand and scarcity of stimulation data on the other. One aspect of such generalizability is across responses to different stimulation frequencies, which is not trivial given that various stimulation frequencies elicit unique inhibitory and excitatory responses^36^. We performed a large-scale combinatorial experiment wherein we trained models for each channel using a dataset with *k* of the 6 stimulation frequencies (including 0Hz, i.e., STIM OFF durations) and validated the models on a test dataset containing all the 6 frequencies. Figure 10 shows the normalized MSE for different values of *k*, averaged across all channels and 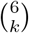 possible models for each value of *k*. We found that using a training dataset with a higher variety of stimulation frequencies proves advantageous in learning a more generalizable model. In particular, the inclusion of 0Hz stimulation (STIM OFF data) in training data resulted in a better model. Given the abundance of such STIM OFF recordings across various datasets, our result indicates that such data can be leveraged to learn more accurate models of stimulation-evoked response models. Additionally, the saturation of the MSE with just 6 frequencies indicates that it is possible to learn a predictive model with a relatively safe and simple experimental design.

Another important aspect of the generalizability is that across temporally-spaced sessions. The RAM dataset consists of multiple stimulation sessions for most subjects, and we assessed whether including training data from a second session can improve modeling accuracy on the first session–as well as a higher bar of whether a model trained only on one session can generalize to a completely unseen session. Per RAM protocols, stimulation sites were varied across sessions. Therefore, generalizability is less expected over STIM ON periods, and we instead tested the hypothesis that constructing a STIM OFF model (without exogenous input) during periods far from any stimulation would exhibit consistency across sessions. If brain dynamics were to be stationarity across different sessions, combining the data directly from these sessions should result in a superior model compared to fitting a model using data from only one session. We thus compared 3 types of switched-linear models over 2 different recording sessions: a model trained on Session 1, a model trained on Session 2, and a third model trained on both sessions. When these models were tested using unseen data from one of the sessions (Session 1), we found the model trained on data only from that session to be the most predictive. This was in fact the case in all 13 subjects. This finding thus indicates the presence of non-stationary or time-varying behavior across sessions. Therefore, models trained over a prior session cannot be directly deployed in a new session, but have to be re-trained continuously to make sure they remain predictive.

## 3 Discussions

In this study, we presented a data-centered framework for modeling the iEEG response induced by intracranial neurostimulation. We highlighted the effectiveness of switched-linear autoregressive models with about 300ms length of history in accurately representing seizure-free iEEG dynamics throughout the pre-STIM to post-STIM duration. Furthermore, we elaborated on efficient methodologies for training such models without requiring supplementary information. We analyzed the models and channels from the standpoint of causal effects of stimulation, both direct and indirect. In addition to the modeling, we experimentally characterized the generalizability of the system identification process across different stimulation frequencies and recording sessions. While this work provides a critical precursor to treating MTL epilepsy via model-based closed-loop neurostimulation, we must highlight that RAM stimulation sites were not located near seizure onset zones (SOZ) and that the data does not contain seizure durations. Nevertheless, the rigorous model-based approach pursued in this paper provides foundational knowledge towards closed-loop neurostimulation in epilepsy, as well as other neurological conditions such as movement disorders and depression.

Our focus on iEEG in this work stems from its importance and gold-standard role as a feedback signal for closed-loop intracranial neurostimulation. In the context of epilepsy, iEEG has long been used for the localization of the SOZ^37, 39, 56^, and subnetworks extracted purely from iEEG data using unsupervised machine learning have been shown to be able to differentiate between SOZ and non-SOZ regions^7, 57^ as well as pre-ictal, onset, and ictal durations^8^. Furthermore, the general predictive modeling methodology used in this study is not restricted to iEEG and can be used, subject to proper modifications, with other forms of neurophysiology and neuroimaging data (see, e.g., Nozari et al.^42^).

A rather undesirable aspect of iEEG recordings in general is the presence of stimulation artifacts^36^ and epileptic discharges. From the perspective of understanding brain function, it is common practice to remove any such effect, often conservatively, and only consider the “clean” data for modeling and analysis^58, 59^. However, when modeling the brain for the purpose of closed-loop detection and control, we need models that accurately capture all the effects and elements that appear in the control loop, including all 3 durations (pre-STIM, STIM ON, post-STIM) and regions (SOZ and non-SOZ, gray and white matter). We can no longer afford to disregard numerous channels and recording durations, as often happens during common iEEG preprocessing. Therefore, we adopt a minimal pre-processing pipeline (cf. Methods) and delegate the task of handling complexities in the data to the data-driven model.

Nonetheless, we investigated the potential effects of artifacts on modeling performance. From Supplementary Figures 7 and 8, we see that the error profile during the 3 durations of pre-stimulation, stimulation, and post-stimulation is dependent on the distance of the target contact from the stimulation site as well as the parameters of the applied stimulation. For channels farther from the anode, resulting in weaker physiologically-evoked response *and* less volume conduction leading to fewer artifacts, the amount of error during the three durations is similar. However, as the total stimulation response (physiological plus artifacts) increases due to less physical distance to the stimulation site and/or larger stimulation amplitude, model prediction errors become significantly larger during STIM ON periods while staying relatively low during the STIM OFF durations. While further research is needed to determine the exact contribution of artifacts to this unexplained variance, this analysis at least ensures that the discussed models have *not* significantly fit to the artifacts and their accuracy is driven by artifact dynamics. Furthermore, since we have carefully time-locked the response time series across repetitions to stimulation onset (Supplementary Figure 2) and neurophysiological evoked responses are typically slower than stim artifacts, it is likely that artifacts are less predictable and therefore contribute more to prediction errors than neurophysiological responses.

An important aspect of our findings is the vast heterogeneity among optimal model structures (let alone parameters) among subjects and even channels within the same subject. This is understandable considering the variations in input effects and the diverse network influences across channels, and only reinforces the need for individualized analysis and treatment in epilepsy^60^. We even observe sufficient heterogeneity among the dynamics of different recording sessions from the same individual to prevent a model tuned to one session from properly generalizing to another. This is conceptually similar to the well-known lack of stability of intracranial multi-unit activity recordings across sessions^61^ but is observed at a significantly larger spatial scale where the physical drift of the recording electrodes relative to the surrounding tissue is less to blame. Nevertheless, across scales, such non-stationarity necessitates adaptive detectors and controllers with constant retuning of models and algorithms^62, 63^.

Our modeling framework is similar to Dynamic Causal Modeling (DCM)^64^ in a number of respects. Bilinear families of dynamic models have a long history in DCM, and switched-linear models with a STIM ON/OFF switching signal are conceptually similar to works that build separate DCM models for STIM ON and STIM OFF periods^44, 65^. Nevertheless, we depart from DCM in a number of ways, including a prediction-error-based parameter estimation methodology^40^ instead of the variational Bayes methodology used in DCM^66^. Furthermore, DCM typically uses *single-lag* bilinear models for fMRI^44^ and complex neural mass models for EEG^67^, whereas we show the power of bilinear autoregressive models *with long input-output histories* in explaining EEG dynamics. These modifications not only make the resulting models more predictively accurate but also significantly more scalable to large-scale recordings. Similar hybrid modeling approaches have also been done for autoregressive models^43^ and provide a broad rationale for the models described in this paper.

In the context of system identification^40^, the experiment design process holds significant importance for and essentially limits the achievable accuracy of any downstream modeling. A good experiment design process involves probing the system with a *persistently exciting* or a diverse set of input frequencies in order to record all the possible dynamical modes of the system^40^. This may even be more important for the brain, as it has been suggested that different stimulation frequencies can result in distinct types of responses, with lower frequencies (10-60 Hz) typically inducing inhibitory effects and higher frequencies (100-200 Hz) producing excitatory responses^36^. Thus, it becomes crucial to incorporate a diverse range of stimulation parameters before modeling. Nevertheless, the conventional approach to experimental design in human studies^41, 43^ often entails repeatedly applying a single, fixed-parameter stimulation. This will allow the models to simply “memorize” the brain’s response to that specific stimulation, without necessarily learning anything generalizable about the brain’s response to neurostimulation. The RAM Parameter Search recordings used in this study are one of very few that encompass a wide range of stimulation frequencies, amplitudes, and durations. However, this dataset is not ideal either, e.g., due to a lack of recordings where changes in stimulation parameters occur without going through a STIM OFF phase. While recent works^34^ have attempted to construct more parameter-rich experiments for non-human primates, similar efforts toward modeling the human brain are largely absent. Nevertheless, the ability to learn predictive models using the RAM dataset, as described in our work, suggests that it is feasible to achieve successful modeling using a relatively small yet carefully chosen set of stimulation parameters.

In our study, we demonstrated a distance-based trend in the MSE_scalar_ − MSE_vector_ plots in Figure 8. Additionally, we showed that electrodes in close proximity to the electrode being modeled play a more significant role in capturing network interactions. However, it is well-known that the brain operates through intricate pathways, and our current approach, which relies on Euclidean distance, may benefit from refinement through more sophisticated methods such as functional or structural connectivity graphs. This could be a promising direction for future research. Additionally, it remains unclear why the MSE_scalar_ − MSE_vector_ trends exhibit substantial variation across different subjects. We investigated electrode coverage and anode location as potential factors influencing these trends, but have not yet identified any statistically significant effects. To develop a robust hypothesis, extending our analysis to include more subjects and sessions will be an essential next step.

**Figure 8:**
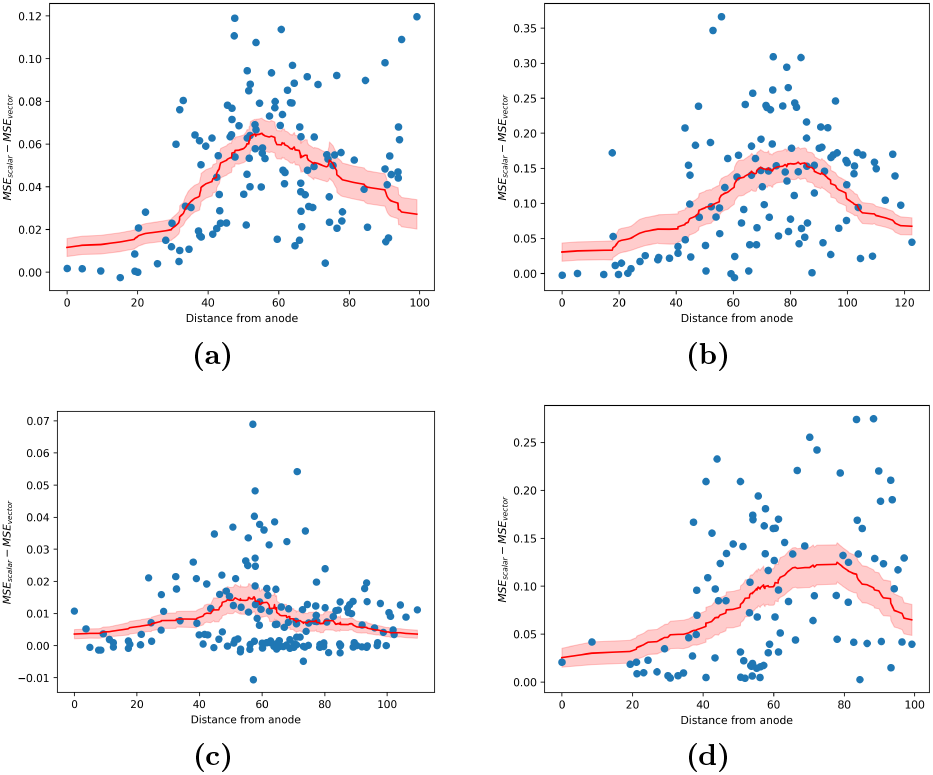
Strength of network interactions as a function of distance from the stimulation site. The strength of causal effects from other nodes (channels) in the network to each node is measured by the MSE advantage MSE_scalar_ − MSE_vector_, where the former (latter) comes from a model in which the history of other nodes’ iEEG values is not (is) used for one-step-ahead prediction. (a)-(d) The MSE advantage for each channel as a function of its distance to the stimulation site (shown in blue) across 4 different subjects. We found that the distance-based moving average of this advantage (shown in red) is initially close to 0, rises with distance until about 70±16 mm, and then decreases thereafter. Plots for the remaining 9 out of 13 subjects have been provided in Supplementary Figure 4

**Figure 9:**
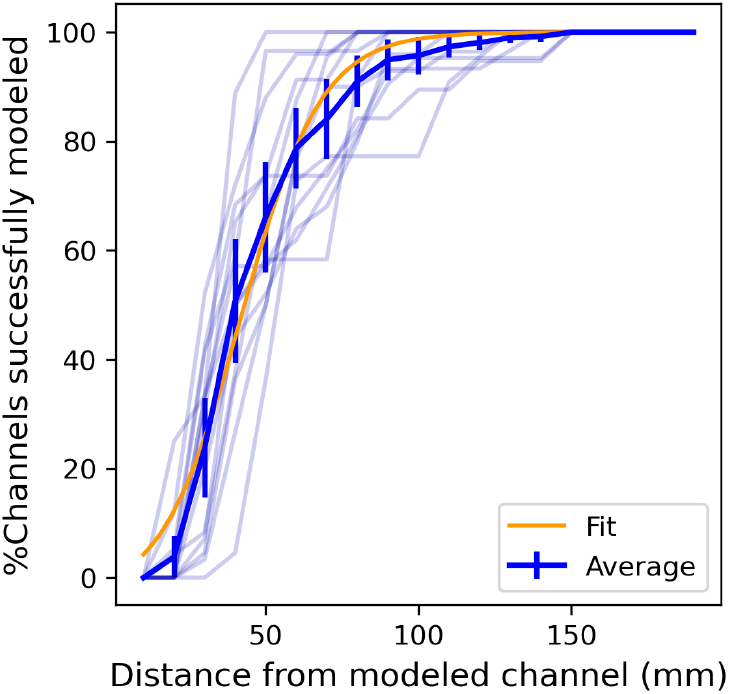
Impact of inter-electrode distance in the modeling of network interactions. For each subject, we plot the % of channels unimpacted by the elimination of an electrode group at a given distance. The build-up within the first 60mm and saturation beyond 80mm indicates that local electrodes are more important in modeling network interactions. Transparent lines illustrate the subject-wise trend, while the opaque line indicates the average trend across all subjects. The orange line shows the best fit sigmoid curve to explain the observed trend.

**Figure 10:**
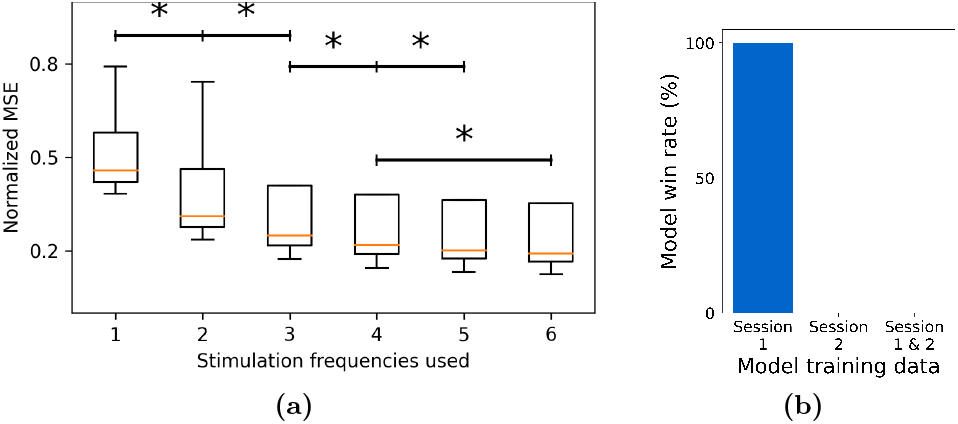
Generalizability of trained models across stimulation frequencies and sessions. (a) We plotted the normalized test MSE as a function of the number of stimulation frequencies used as part of the training set. We found that including more frequencies results in a more generalizable model. (b) However, we did not find it to be beneficial when we extend the dataset using recordings from multiple sessions. Specifically, when we considered 2 sessions and the 3 possible models that can be trained from them (using only session 1 or only session 2 or both in the training set), we found that unseen data from session 1 was best predicted by the model trained on data from Session 1 only. (\**p* < 0.05)

Future steps towards the development of a complete model-based closed-loop seizure control system require even richer and more extensive datasets that encompass simultaneous seizure and neurostimulation, enabling models to learn the interactions between ictal and stimulation-evoked dynamics. Such data, however, is challenging to collect for a number of reasons. Despite ongoing advancements in seizure detection, accurately and reliably timing stimulation relative to seizure onset remains a major challenge, which is further exacerbated by the heterogeneity of seizure events even within the same individual. Further, iEEG recordings have significantly better spatial and temporal coverage and algorithmic flexibility during acute seizure monitoring, but manipulating intrinsic seizure dynamics can interfere with medical diagnostics at this stage. Animal and computational models can therefore play an invaluable role before algorithmic designs can be translated to humans.

## 4 Methods

### Data

Throughout this work we use data from the Parameter Search experiments of the Restoring Active Memory (RAM) project, consisting of multichannel resting state, task-induced, and neurostimulation-induced iEEG responses of over 80 human subjects^68^. As reported extensively in the RAM documentation, the highest stimulation amplitude (*A*_safe_) that could be applied to each subject without resulting in after discharges were first measured. This value was then used in the PS2 experiments which we use in this work, providing subjects with 500ms fixed-duration stimulations of biphasic pulse trains delivered at varying combinations of current amplitudes (3 levels: *A*_safe_, *A*_safe_ − 0.25mA and *A*_safe_ − 0.5mA) and frequencies (single pulse, 10Hz, 25Hz, 50Hz, 100Hz, or 200Hz). Each subject has undergone multiple stimulation sessions, ranging from 2 to 5, with each session involving a different stimulus (anode-cathode) location as shown in Figure 1.

During each session, pulse trains of each combination of amplitudes and frequencies were repeatedly delivered multiple times (about 20-30 times, equivalent to 30 minutes of total experimentation). Such exhaustive repeated parameter sweep data is rare in stimulation-evoked iEEG and makes this dataset particularly rich for the modeling purposes of this study. To partly control for the great variability among subjects and the high computational cost of numerical experiments, we limited our analyses to subjects with an iEEG sampling frequency of 1000Hz. Additionally, we excluded subjects with stimulation outside the medial temporal lobe (MTL) were removed, resulting in 13 subjects for further analysis.

### Preprocessing and data decomposition

As noted earlier (see Discussions), we used minimal preprocessing in order to build models that can capture stimulation-evoked iEEG dynamics in its full complexity. We removed power line harmonics by applying 4th-order Butterworth notch filters at 60Hz, 120Hz, and 180Hz. We then detrended the signal using a first-order polynomial fit to remove linear drifts and z-scored the data to standardize signals across all subjects and channels. Finally, we selected windows of length 1500ms around each stimulation. These consisted of 500ms pre-stim, 500ms stim, and 500ms post-stim for pulse train stimulations and 750ms prestim and 750ms post-stim for single-pulse stimulations. This was due to the fact that only about 1/6 of the recordings corresponded to durations where stimulation was applied. Hence, using the full recordings could (and did, based on our pilot analyses) bias the models towards fitting STIM OFF durations and ignoring evoked responses.

We define the input time-series *u*(·) based on the stimulation timing and parameters. If a pulse train with frequency *f* and amplitude *a* was applied starting from time *t*_*k*_ (always lasting for 500ms), then we set *u*(*t*_*k*_ + Δ*t*) = *a* for all Δ*t* ∈ [0, 500] that are integer multiples of 1000*/f* and *u*(*t*) = 0 otherwise. The modulating term 𝒰(·) in Eq. (1) was set as 𝒰(*t*) = *a* (amplitude-weighted switched linear) or 𝒰(*t*) = 1 (switched linear) for all *t* within stimulation durations [*t*_*k*_, *t*_*k*_ + 500], and 𝒰(*t*) = 0 for other duration. In the case of the complete (fully) bilinear model, we set 𝒰 = *u*, which can be thought of as a more natural choice for modulation. See Supplementary Figure 1 for further clarification.

The input times recorded in the RAM data were frequently found to be inconsistent with the actual times of stimulation delivery, as indicated by the delays in the output response of the anode channel (Supplementary Figure 2). To address this discrepancy, we identified the actual time of input by thresholding the difference Δ*y*(*t*) = *y*(*t*) − *y*(*t* − 1) in the anode channel. This allowed us to detect the precise time of stimulation delivery, and we used this onset time for time-locking instead of the reported onset time in the dataset.

Unless otherwise stated, we split the processed iEEG recordings into training and test as follows. Each 1500ms window (corresponding to each instance of stimulation, see above) was split into 5 continuous folds of data (300ms each), out of which 4 randomly selected folds were used for training, and the remaining fold was used for testing.

### Models

The main focus of our work is the dynamic model described in Eq. (1), which is used to make one-step ahead predictions about the iEEG response given past iEEG and input history lags as regression features. We denote the iEEG history of length *L* for channel *k* at time *t* as the vector 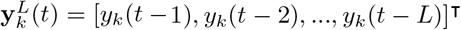. Similarly, the input lags vector **u**^*M*^ (*t*) of length *M* is given by [*u*(*t* − 1), *u*(*t* − 2), …, *u*(*t* − *M*)]^⊺^.

Several families of models used in this work are special cases of the model in Eq. (1). A linear autoregressive model is obtained by setting the matrices **b**_*k*_, **c**_*K*_, and **d**_*k,i*_ to 0. The learnable (not necessarily zero) parameter sets of the subclasses of Eq. (1) used in this work are listed in Table 1.

**Table 1:** Parameter sets of linear and bilinear/switched linear (BL/SL) models used in our work. Note that bilinear and switched linear models have the same set of nonzero weights and only differ in their encoding of 𝒰(see ‘Preprocessing and data decomposition’ above). In each case, the shown matrices are learned from data while matrices outside of this set are assigned to 0.

| Model | Parameters $\theta$ |
| --- | --- |
| AR | $\{\mathbf{a}_k\}$ |
| ARX | $\{\mathbf{a}_k, \mathbf{b}_k\}$ |
| VAR | $\{\mathbf{a}_k, \mathbf{d}_{k,i}\}$ |
| VARX | $\{\mathbf{a}_k, \mathbf{b}_k, \mathbf{d}_{k,i}\}$ |
| BL/SL AR | $\{\mathbf{a}_k, \mathbf{c}_k\}$ |
| BL/SL ARX | $\{\mathbf{a}_k, \mathbf{b}_k, \mathbf{c}_k\}$ |
| BL/SL VAR | $\{\mathbf{a}_k, \mathbf{c}_k, \mathbf{d}_{k,i}\}$ |
| BL/SL VARX | $\{\mathbf{a}_k, \mathbf{b}_k, \mathbf{c}_k, \mathbf{d}_{k,i}\}$ |

Determining the structure of Eq. (1) involves choosing appropriate model orders (*L, M*, and *P*) as well as determining the subset of {**a**_*k*_, **b**_*k*_, **c**_*k*_, **d**_*k,i*_} that can be nonzero. We make all such choices in a data-driven manner. This is particularly important when we want to fit personalized models, where we need to account for the knowingly large subject-to-subject variability^60, 69^. Given the compound and combinatorial nature of structural choices, we also use a hierarchical bottom-up approach where we tune low-level quantities (e.g., number of autoregressive lags) before selecting higher-level choices (e.g., type of nonlinearity or presence of network interactions). For each channel, we first assume an ARX model and find the pair of iEEG and input lags (i.e., *L* and *M*) that maximizes the model accuracy. Fixing the obtained values of (*L, M*), we train the models described in Table 1 while keeping the value of the maximum network lags *P* as 1. Once we statistically determine the best model structure (i.e., linear or bilinear/switched), we use it to determine the optimal value of *P* by evaluating the validation MSE of models fit with different values of *P* (increased from 1 to 100 in steps of 10).

In addition to the above models, we trained other nonlinear models, as follows:

*Artificial Neural Network (ANN)*: A 2-layer feed-forward network with ReLU activation, with 500 and 50 nodes in the first and second layer respectively, and inputs being 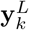 and **u**^*M*^. We opted for the ReLU activation function due to our observation that it yielded superior performance compared to alternative nonlinear functions, such as tanh.

*Long Short-Term Network (LSTM)*: Standard single layer LSTM with *y*_*k*_(*t* − 1) and *u*(*t* − 1) as input and expected output Δ*y*_*k*_(*t*)

*Sparse Identification of Nonlinear Dynamics (SINDy)* using 3^*rd*^ degree polynomial expansion *Nonlinear ARX (NLARX)* specified in MATLAB^70^, model order (50, 50, 1) used for all channels

### Learning

We train the free parameters of each model by minimizing the regression mean squared error

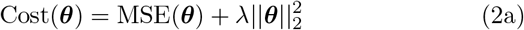

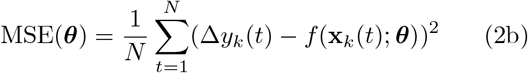

where *f* (·) is any of the aforementioned parametric families of models and **x**_*k*_(*t*) denotes the regression features of the *t*^*th*^ sample and the *k*^*th*^ iEEG channel. This may include the lags of iEEG recording of the *k*^*th*^ channel 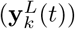, that of other channels 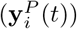, and input lags (**u**^*M*^ (*t*)). This minimization is *l*_2_-regularized to avoid overfitting. Specifically, we added a cost of 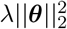 to the MSE cost function, where *λ* is the penalty factor and was empirically fixed to a value of 0.1.

### Performance metric

We evaluated trained models over test data based on a number of metrics.One metric is test (normalized) MSE which is also computed from Eq. (2) but computed over test data. Sample durations with different levels of NMSE have been provided in Supplementary Figure 3. Since MSE (as with any mean-based measure) can be susceptible to outliers, we also compared the models based on a one-sided Wilcoxon signed-rank test with Bonferroni correction for comparisons between multiple pairs. Once we determine the best models for each channel across all the subjects, we define the model *win rate* for any model *f*_*i*_ as

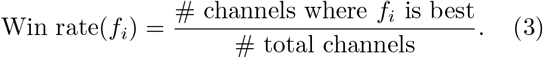

### Sparse ARX modeling via delay embedding

When comparing dense vs. sparse autoregressive models, we compared a regular ARX model (cf. Table 1) with a sparse version

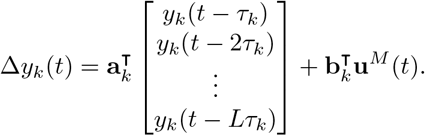

The tuning of the delay time *τ*_*k*_ has been studied extensively in the delay embedding literature^49^. In this work, we select a separate optimal *τ*_*k*_ for each subject and channel using the auto embedding method described in Otani et al.^51^.

### Indirect test of linearity

The results demonstrated in Figure 4 were obtained as follows. For each channel, we divided the iEEG dataset into 3 sub-datasets, with the first one including only STIM OFF data while the other 2 contained only STIM ON durations. Using the first dataset, we trained an AR model with parameter matrices 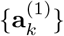. The second dataset was used to learn an ARX model with parameter matrices 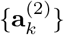 and 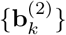. Finally, a second ARX model was trained on dataset 2 by fixing autoregressive weights to 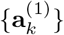 and only learning 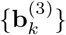.

We then tested the two ARX models 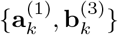 and 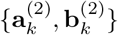 on the third dataset. If the underlying process is indeed linear, we expect the autoregressive weights ({**a**_*k*_}) to remain the same during STIM ON and STIM OFF durations, which would result in the ARX model 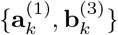 performing as good as 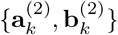. However, if the process is nonlinear, the latter would perform significantly better because of having both {**a**_*k*_} and {**b**_*k*_} trained on STIM ON data.

### Bootstrap Test

For each subject and channel, we used a bootstrap test to validate if the presence of input lags significantly improved the prediction accuracy (i.e., whether a direct causal effect from input to that channel existed). From the original dataset 𝒟 = {*Y*_*k*_, *U, Y*_−*k*_, Δ*y*_*k*_} for channel *k* (where *Y*_−*k*_ corresponds to the iEEG information from all channels other than *k*) we created a synthetic one 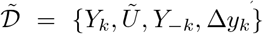 by randomly shuffling the input history data *U* (·) across time samples (i.e, 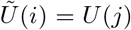 for some random *j*). We then split this dataset into training and test sets, learn a dynamical model, and compute the corresponding test MSE as before. We repeated this process 100 times and compared the test MSE of the unshuffled model to the distribution of test MSEs from shuffled models to compute a sample-based *p*-value. We reject the null hypothesis of no input effect (i.e., conclude that a direct causal effect from input to that channel existed) if *p* < 0.05.

In addition to input effects, we also validated the causal effects of network interactions in each subject-channel using a similar bootstrap test, wherein we generate synthetic datasets for each channel by randomly shuffling the iEEG past lags of other available channels. Specifically, a synthetic dataset 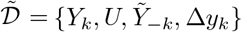 was created by randomly shuffling the network information history data *Y*_−*k*_(·) across time samples (i.e, 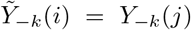 for some random *j*). The test for network effects follows the same experimental structure as described in the previous paragraph.

### Learning models on subsets of stimulation frequencies

The results reported in Figure 10 were generated as follows. We first split the dataset for each subject based on the stimulation frequency during each window and created multiple training sets corresponding to all subsets of {0 (*no stim*), 10, 25, 50, 100, 200}. For each subset of frequencies, we fit a switched ARX model and tested it on the complementary subset of frequencies omitted in the training set. The normalized MSEs for all the combinations are reported in Figure 10.

### Statistics and Reproducibility

Statistical testing were performed using the *stats* API of the *SciPy* library in Python. Model comparisons were performed using the one-sided Wilcoxon signed-rank test and paired *t* -test. We have used Bonferroni correction to account for multiple comparisons. Significance was declared when we obtained *p* < 0.05. We ensure reproducibility of our results (e.g., train-test splits, bootstrapping) by fixing the *NumPy* ‘s random number generator.

## Acknowledgements

The research performed in this study was supported in part by National Science Foundation Award Number 2239654.

## Author Contributions

EN designed and supervised the study; GA performed the analyses; KAD co-supervised the selection and processing of the data and the empirical interpretation of the results; all authors wrote the manuscript.

## Data availability

The iEEG data used in this study is publicly available from the RAM Public Data Release at http://memory.psych.upenn.edu/RAM. The numerical source data corresponding to the generated plots can be accessed from https://doi.org/10.6084/m9.figshare.26333371.v1.

## Code availability

Python code for the analyses in this study is publicly available at https://github.com/nozarilab/2023Acharya_evoked_ieeg.

