## Supplementary figures for "Predictive Modeling of Evoked Intracranial EEG Response to Medial Temporal Lobe Stimulation in Patients with Epilepsy"

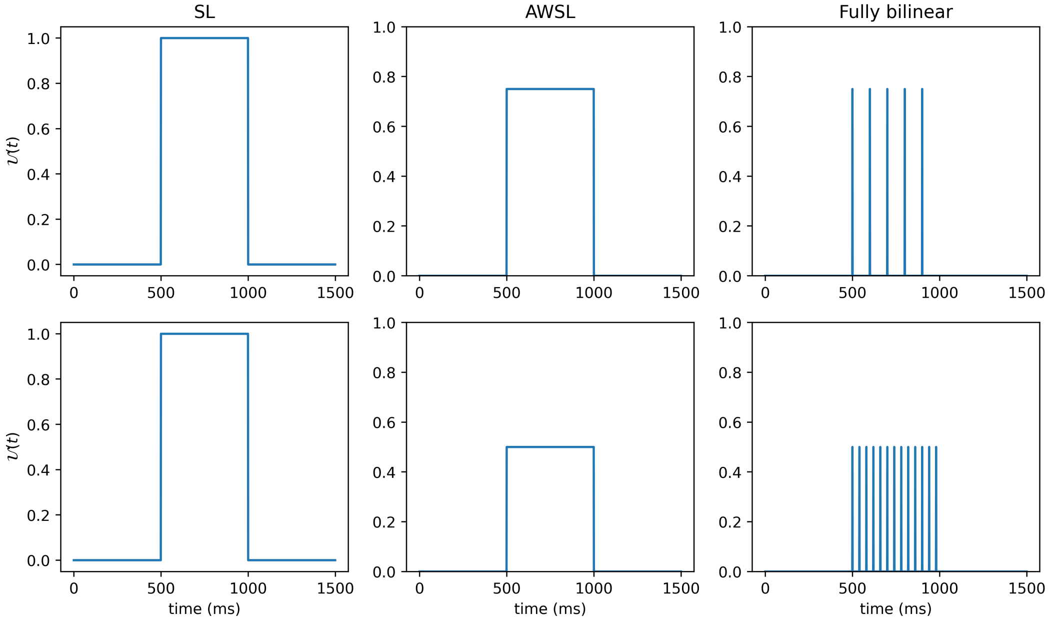


Supplementary Figure 1: $\mathcal{U}$ **for different types of bilinear models mentioned in our work.** The first row corresponds to a duration associated with a $750\mu A$ current and 10Hz stimulation. In the switched-linear (SL) model, $\mathcal{U}(t)$ is 1 during the 500ms STIM-ON period and 0 elsewhere. In contrast, in the amplitude-weighted SL model (AWSL), $\mathcal{U}(t)$ equals the applied amplitude ($0.750mA$ in this case) during the STIM-ON period. In the fully bilinear model, $\mathcal{U}(t)$ matches the input signal $u(t)$, displaying the individual instances of stimulation separated temporally based on the stimulation frequency (100ms in this case). The second row shows the same for an input with a $500\mu A$ current and 25Hz stimulation.

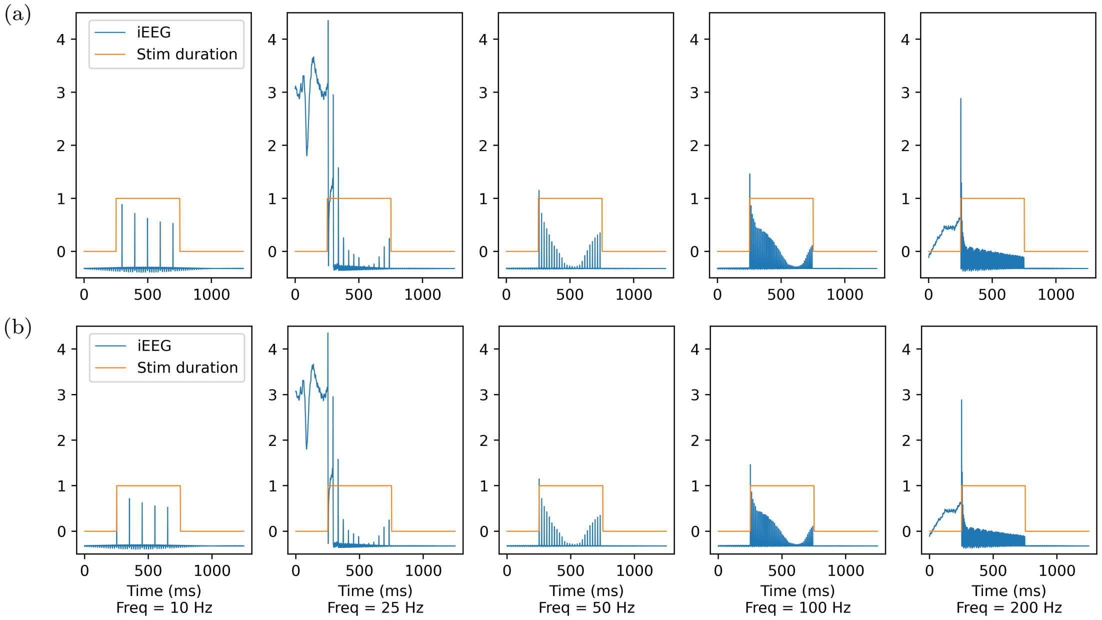


Supplementary Figure 2: **Correcting input-output phase mismatch.** The input times provided in the data generally may not reflect the precise time of stimulation delivery, as suggested by the delay in the output response of the anode channel. This mismatch is particularly prominent for durations associated with the 10Hz stimulation frequency. To address this, we identified the actual time of input by thresholding the difference $\Delta y(t) = y(t) - y(t-1)$ in the anode channel, detecting the precise time of stimulation delivery, and using this onset time for time-locking instead of the reported onset time in the dataset. The figure shows iEEG data with both the uncorrected input series and the corrected input series for different stimulation frequencies, highlighting the improvement in temporal alignment between the input and output signals.

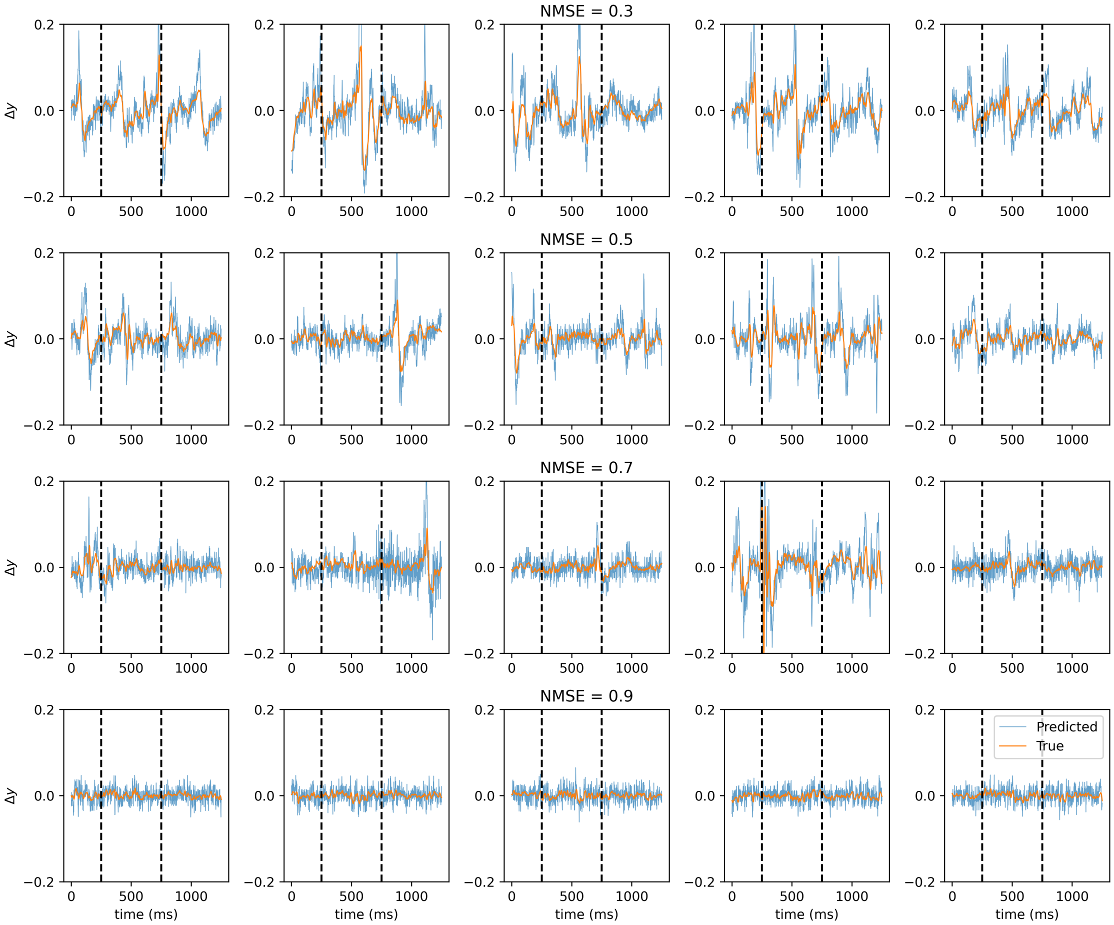


Supplementary Figure 3: **Sample durations of predicted vs. actual iEEG response for different levels of normalized mean squared error (NMSE).** Each plot shows the true (in blue) and predicted (in orange) values of $\Delta y(t) = y(t) - y(t-1)$ as a function of time (cf. Eq. (1) in the main text). Within each plot, the duration within the vertical grid lines corresponds to the STIM-ON period. Each row corresponds to durations where the NMSE is in the range $[a, a + 0.1]$, with $a$ increasing from 0.3 to 0.9 from the first to the last row.

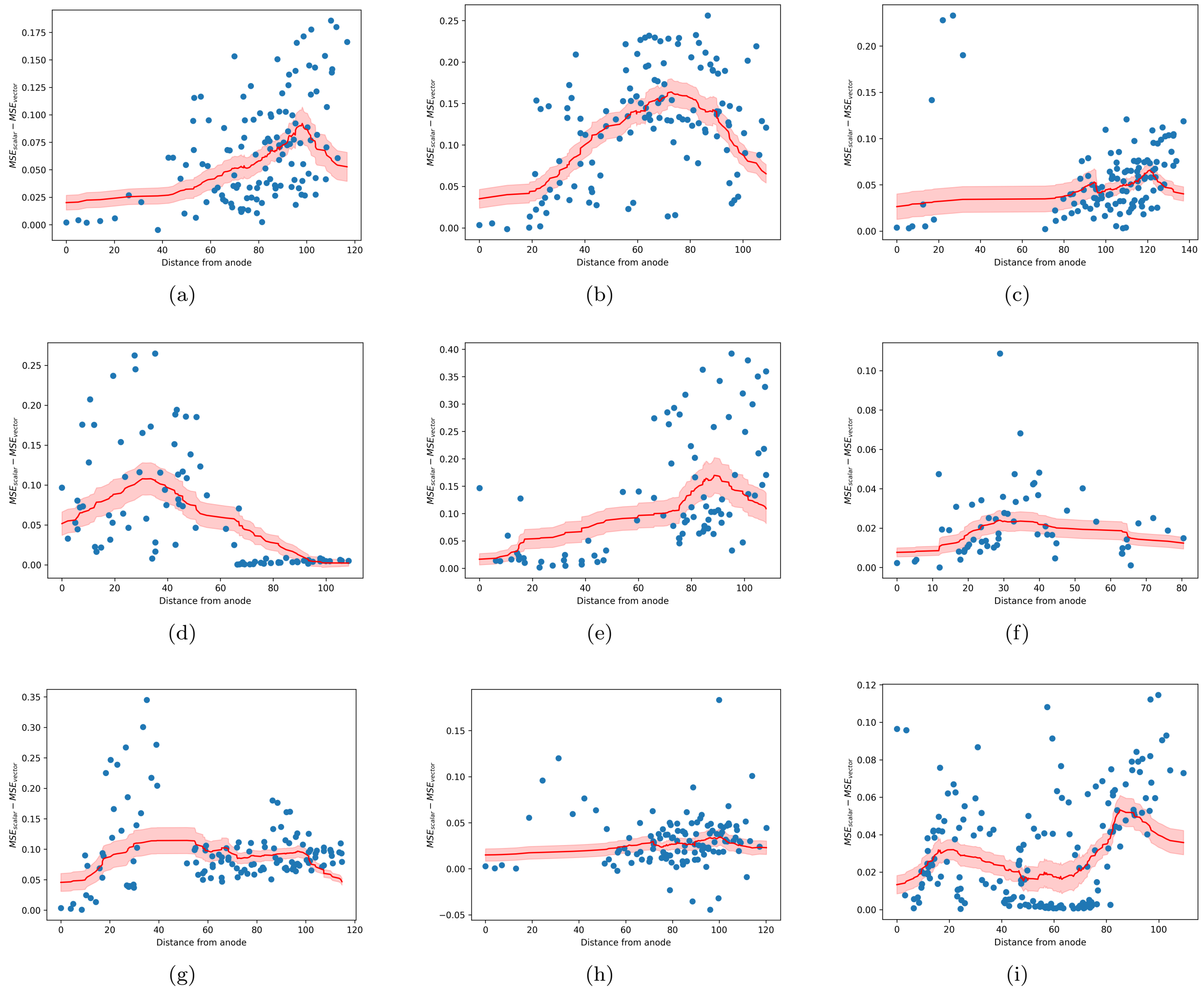


Supplementary Figure 4: **Distance-based network interaction trend plots for the 9 subjects not shown in Main Figure 8**. Consistent trends are observed, with elevated strength for electrodes that are mid-range (neither too close to nor too far from) the anode. Significant heterogeneity is nevertheless also observed among subjects, including heterogeneity in the distance at which the peak occurs.

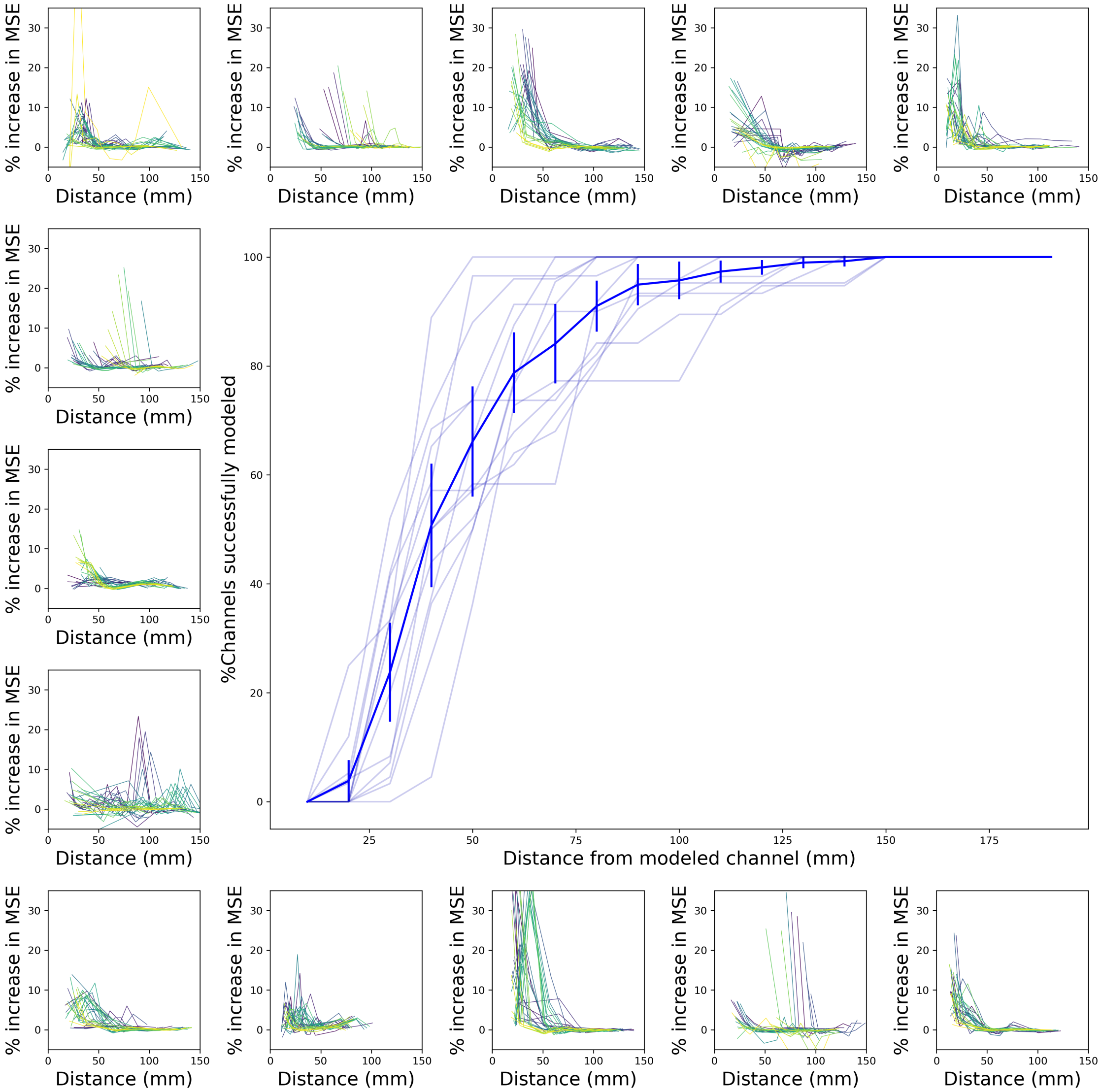


Supplementary Figure 5: **Backward elimination of features reveals that local electrodes account for** 98% **of network dynamics.** For each subject-channel, we identified 10 groups of channels, each consisting of $n_g = 10\%$ of the total number of channels ($= C$), and sorted them based on their proximity to the target channel (the channel being modeled). Subsequently, we construct switched VARX models by iteratively excluding the features associated with each electrode-group in a leave-one-out fashion. To elaborate, we fixed the weights $D_k^i$ (cf. Eq. (1)) associated with the last $n_g$ channels within a distance $d$ mm to zero and learned the remaining weights using least squares. We then noted the increase in test MSE in comparison to a model with all $C$ channels. This process was repeated for all groups and channels, and we plotted the performance degradation (Y-axis: % increase in MSE) as a function of group-distance (X-axis) in a subject-wise manner as represented by each of the small subpanels (different line colors in each subpanel correspond to various channels within the same subject). For most channels across subjects, we find that the performance drop by removing the closest electrode-group is typically over 10%. Further, removing the farthest group barely affects the modeling, indicating a preference for local interactions. In the big panel, we plot for each subject the percentage of channels successfully modeled (Y-axis:% channels where % MSE increase as a result of eliminating a group is $< 2\%$) as a function of the distance of the eliminated group (X-axis). Transparent lines in this plot illustrate the subject-wise trend, while the opaque line indicates the average trend across all subjects. We see that for approximately $50 \pm 12\%$ of the channels, it is sufficient to include just the local electrodes within a 40mm radius of the modeled channel to yield a performance *equivalent to* (i.e., less than %2 different from) using all $C$ channels. This proportion rises to $80 \pm 9\%$ of the channels when the radius is increased to 60mm. These results suggest a possibility of modeling iEEG response at a local level and developing models with sparser regression features, all of which are critical for future implementations in chronic implants.

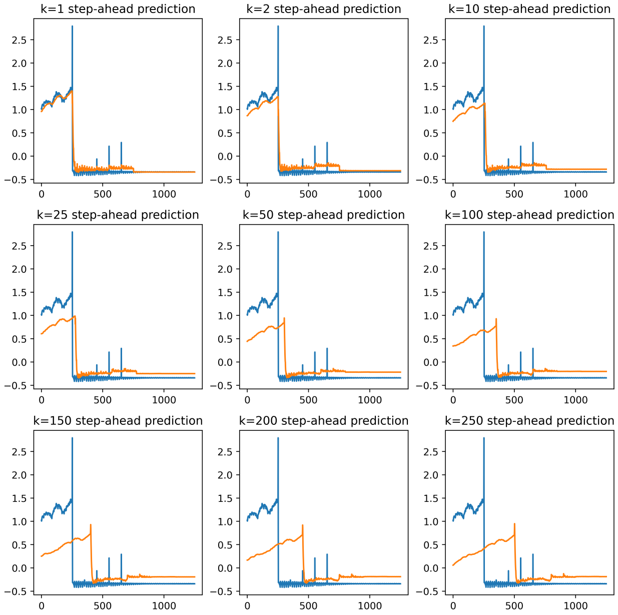


(a)

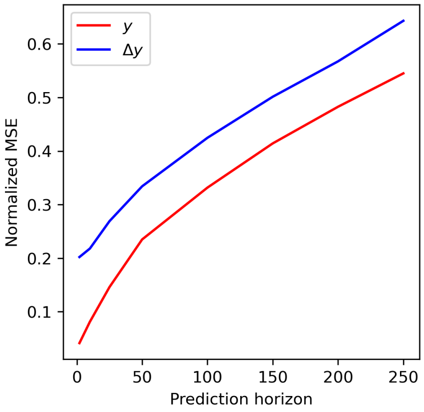


(b)

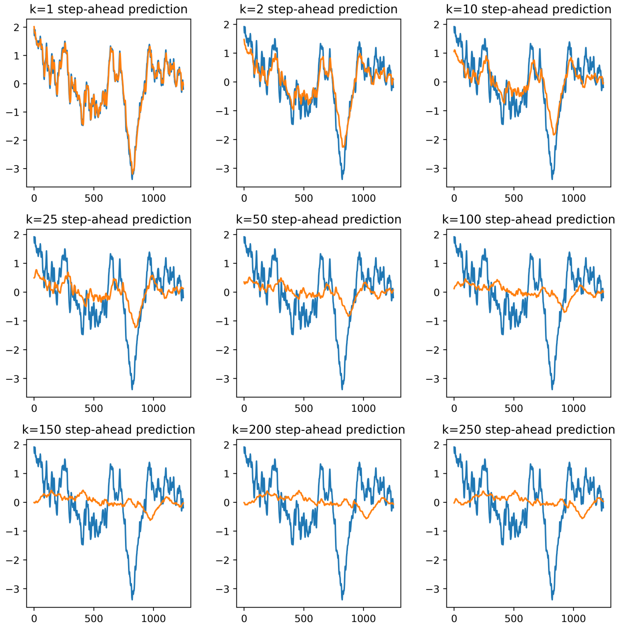


(c)

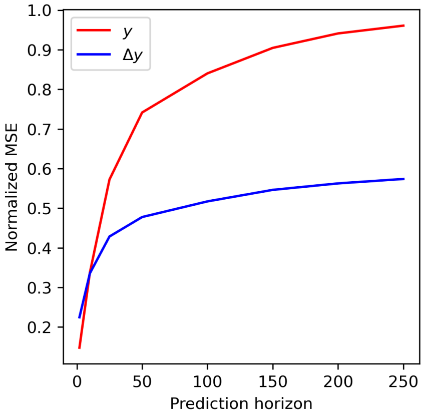


(d)

Supplementary Figure 6: **Effect of increasing prediction horizon on modeling accuracy. (a)** Each subplot shows $y(t)$ and $\hat{y}(t|t-k)$ for the anode channel of a randomly selected subject (Subject: R1117J) and various prediction horizons $k$. For each $k$, $\hat{y}(t|t-k)$ is the prediction of $y(t)$ given past values of $u(\tau)$ and $y(\tau)$ up to time $\tau = t-k$ modeled using the optimized Switched-Linear model. **(b)** Mean normalized MSE of $y(t)$ and $\Delta y(t)$ as a function of prediction horizon $k$ for the same subject and channel in (a). **(c,d)** Similar to (a,b) but for a randomly selected channel with low stimulation artifact in the same subject. As expected, the 1-step ahead predictions closely align with the original $y(t)$. However, as $k$ increases to 250, the forecasts become progressively less accurate, with the fit at larger $k$ values resembling a delayed low-pass filter. This trend is consistently observed across all iEEG channels and durations.

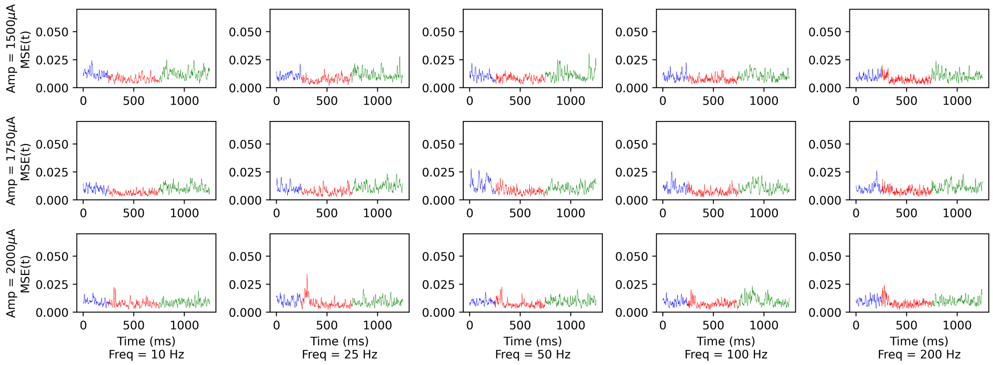


(a)

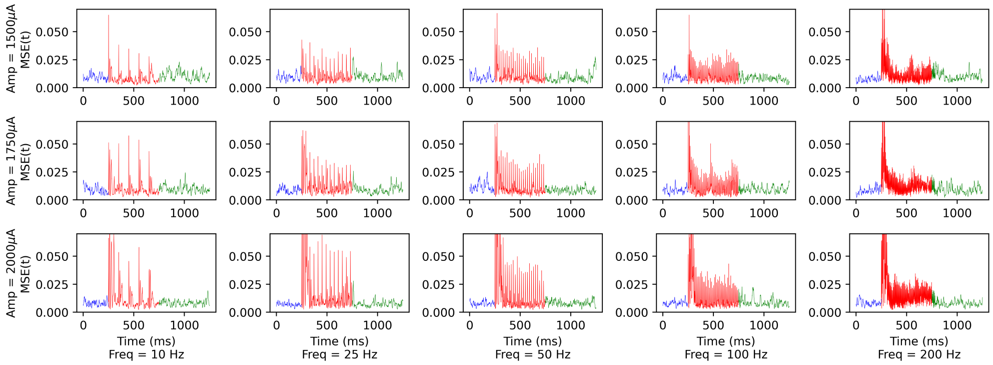


(b)

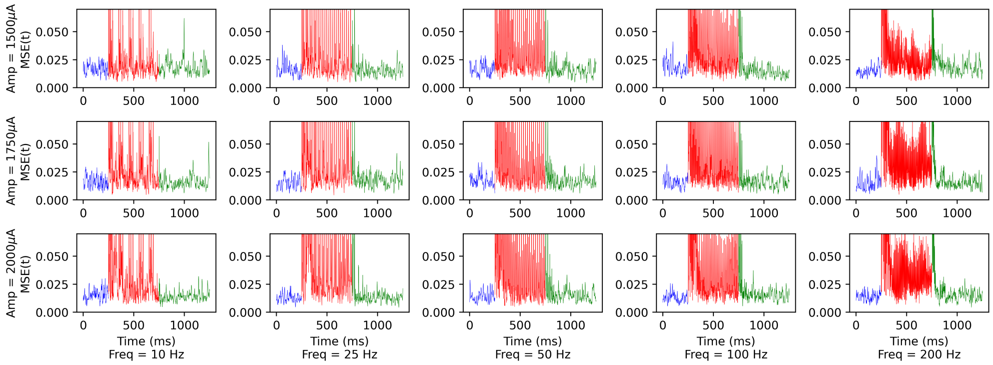


(c)

Supplementary Figure 7: **Effects of stimulation artifacts on modeling performance (Subject ID: R1136N). (a)** The cross-validated (test) performance of the optimal switched-linear model for a randomly selected subject (R1136N) and a target electrode that is far from the stimulation site (60mm). Stimulation artifacts are minimal due to the large distance between the target and stimulated channels. The 3-row by 5-column array of subplots represents different combinations of stimulation amplitudes and frequencies. Within each subplot, the average of the squared prediction error time series over all 24 repetitions of that (amplitude, frequency) combination is depicted in blue for the pre-stimulation duration, in red for the 500 ms stimulation duration, and in green for the post-stimulation duration. **(b, c)** Same as in (a) but for target electrodes that have a medium (42mm) and near (31mm) distance from the stimulation site, respectively. In panel (a) where the stimulation artifacts are minimal, the prediction error levels for the three durations are comparable. As the artifact level increases in panels (b) and (c), the prediction errors during the stimulation period become significantly higher than those during the pre-stimulation and post-stimulation periods.

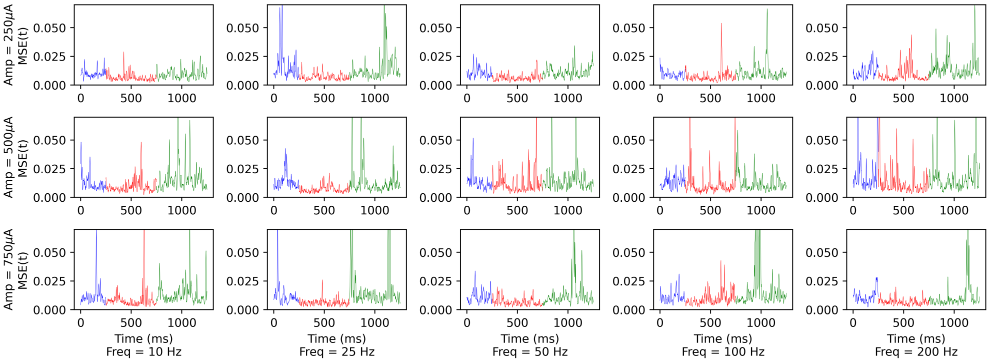


(a)

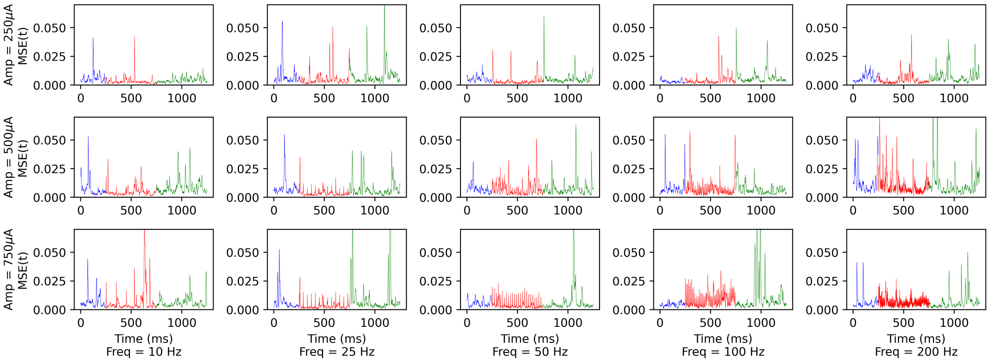


(b)

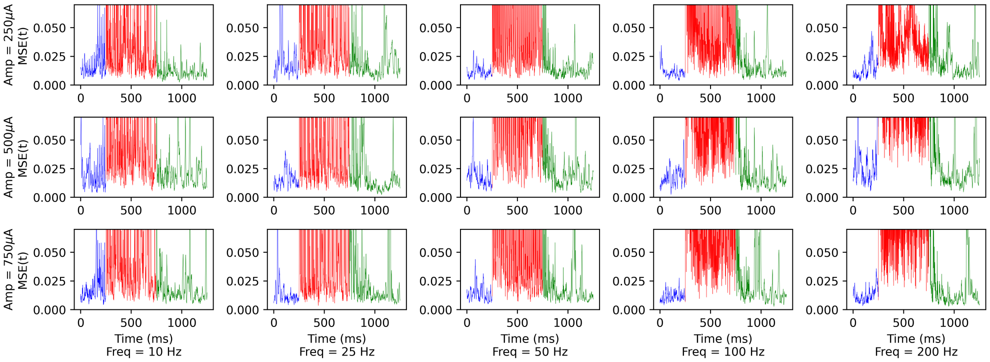


(c)

Supplementary Figure 8: **Effects of stimulation artifacts on modeling performance (Subject ID: R1094T).** Panels parallel those in Supplementary Figure 7 for a different subject (R1094T). The target electrodes in panels (a-c) are within 124mm, 92mm, and 14 mm from the stimulation site, respectively.
